# Paradise fish (*Macropodus opercularis*) as a novel translational model for emotional and cognitive function

**DOI:** 10.1101/2024.09.06.611721

**Authors:** Zoltán K. Varga, Diána Pejtsik, Tímea Csorvás, Éva Mikics, Ádám Miklósi, Máté Varga

## Abstract

Zebrafish have revolutionised physiological screening in vertebrates, but the interpretation of individual-based behavioural assays is burdened by the strong sociality of this species. We propose the use of a solitaire fish species, the paradise fish to keep the advantages and compensate for the limitations of the zebrafish model. We compared juvenile paradise fish and zebrafish in social and non-social exploratory tasks, anxiety tests and in a working memory assay to assess their performance in these individual-based models. We found that in contrast to zebrafish, paradise fish did not show social approach in the U-shape sociability test, their novelty exploration was not biased by the presence of a conspecific in the slalom test, and was not impaired by social isolation in the showjump or in the swimming plus-maze (SPM) anxiety tests. While social circumstances did not affect the anxiety of paradise fish, it was sensitive to the anxiolytic drug buspirone. Intra- and interest repeatability measures of the anxiety tests revealed that paradise fish express more consistent exploratory and defensive behaviours regarding time and context compared to zebrafish. Behavioural consistency in paradise fish was also supported by arm alternation as a predominant choice of exploration in the y-maze task. In summary, our results indicate that the behaviour of paradise fish is less biased by social cues and is more effective and repeatable in individual-based assays than zebrafish behaviour. We hypothesise that the two phenomena are connected and novelty exploration, anxiety and working memory can be more reliably measured and are translatable in a solitaire species.

## INTRODUCTION

Animal models are indispensable for biological research, especially to understand the mechanisms and causative relationships of biological processes. Different fields have benefited from a wide range of model species, each bringing with them specific trade-offs between simplicity versus similarity to humans^1–5^. Fishes are positioned towards the middle of such imaginary scales, manifesting the hallmark features of a conserved vertebrate body plan, yet in a relatively simple form. While over the past decades different scientific fields favored different fish species^6–9^, zebrafish (*Danio rerio*) have lately become the predominant model-of-choice for biomedical sciences^10,11^. The popularity of zebrafish, however, is partly due to serendipity: it became one of the dominant models of genetics and as genetic tools started to dominate life sciences, zebrafish research found new niches.

Nowadays high-throughput and high-resolution screening can be combined with an ever expanding genetic toolkit, making zebrafish an extremely powerful model species, almost unparalleled among other vertebrates^12–14^. Nowhere else is this more apparent than in neuroscience: fish models let us assess neuronal activity simultaneously and continuously in the whole brain and on the level of single cells, in living animals^15^. These *in vivo* imaging methods combined with cell-specific manipulation techniques will elevate our knowledge about brain mechanism and function to a whole new level.

Despite these advances, the validity and generalisability of zebrafish behaviour is still a missing link in this promising model system. In their natural habitat zebrafish only exist in shoals^16^. Their behavioural repertoire has evolved to live in such a simple social structure and their individual behaviour is determined and only interpretable in such circumstances. Internal states and behavioural responses of zebrafish are highly affected by either the presence or the absence of conspecifics^17–24^. Social buffering and social contagion mechanisms homogenise the behaviour of a group and from that point the group itself should be considered as an independent sample^22,23^. Also, group-level assessment of behaviour cannot be correlated with individual physiological data. If individual-level behaviour is assessed by separating zebrafish individuals from their shoal, that will constitute a major and specific perturbation on the external and internal state of the given individuals^24–27^. Either way, in tests that primarily aim to measure emotional or cognitive functions the behavior of the observed individuals will be at least partially biased. This begets the question of what exactly we can learn from the performance of an isolated individual belonging to a species which evolved to perform in a group?

Here, we propose the use of an alternative fish species, the paradise fish (*Macropodus opercularis*) to keep the advantages and compensate for the limitations of the zebrafish model. A reference genome and details of the effective housing of paradise fish were currently published^28,29^, while its behavioural repertoire was already well-described, including complex defensive^30–32^, social^33,34^ and cognitive responses^35,36^. Paradise fish individuals live in a less cohesive social structure than zebrafish, one that resembles more human social settings. Paradise fish show context-dependent social activity including complex courting, mating, aggressive or parental behaviour, but mostly function as soliters. We hypothesise that the behaviour of a fish that naturally copes with challenges alone has more intact and hence more reliable responses when facing different tasks alone. In this study, we aimed to compare the performance of juvenile zebrafish and paradise fish in individual-based behavioural assays and to determine the biasing effects of their social environment on their performance. We assessed i) the sociability of the two species, with or without the presence of the other species, and ii) the effect of conspecifics on exploratory behaviour and anxiety. Furthermore, we compared the performance of individual zebrafish and paradise fish in tasks modelling higher order processes, such as iii) responses to anxiolytics, iv) behavioural consistency and v) working memory. We found that compared to zebrafish paradise fish are less biased by extrinsic social and non-social factors and show more consistent and efficient performance in paradigms that model human phenomena. We conclude that paradise fish is a more reliable model to be used in individual-based assays while zebrafish should be the model of choice in tasks that involve social challenges.

## RESULTS

In the first three experiments we aimed to compare how the presence or absence of a conspecific affects the behaviour of the two species.

### Paradise fish show less direct social interactions compared to zebrafish

In our first experiment, we aimed to compare the sociability of zebrafish and paradise fish in the U-shaped sociability test^37^ in baseline or perturbed conditions. As a baseline, we measured the preference towards a single stimulus conspecific (intra-species challenge), while in perturbed conditions the other species were also presented as stimulus (double-species challenge). As a control we also measured the preference towards the other species per se (inter-species challenge) (Figure 1A).

**Figure 1.**
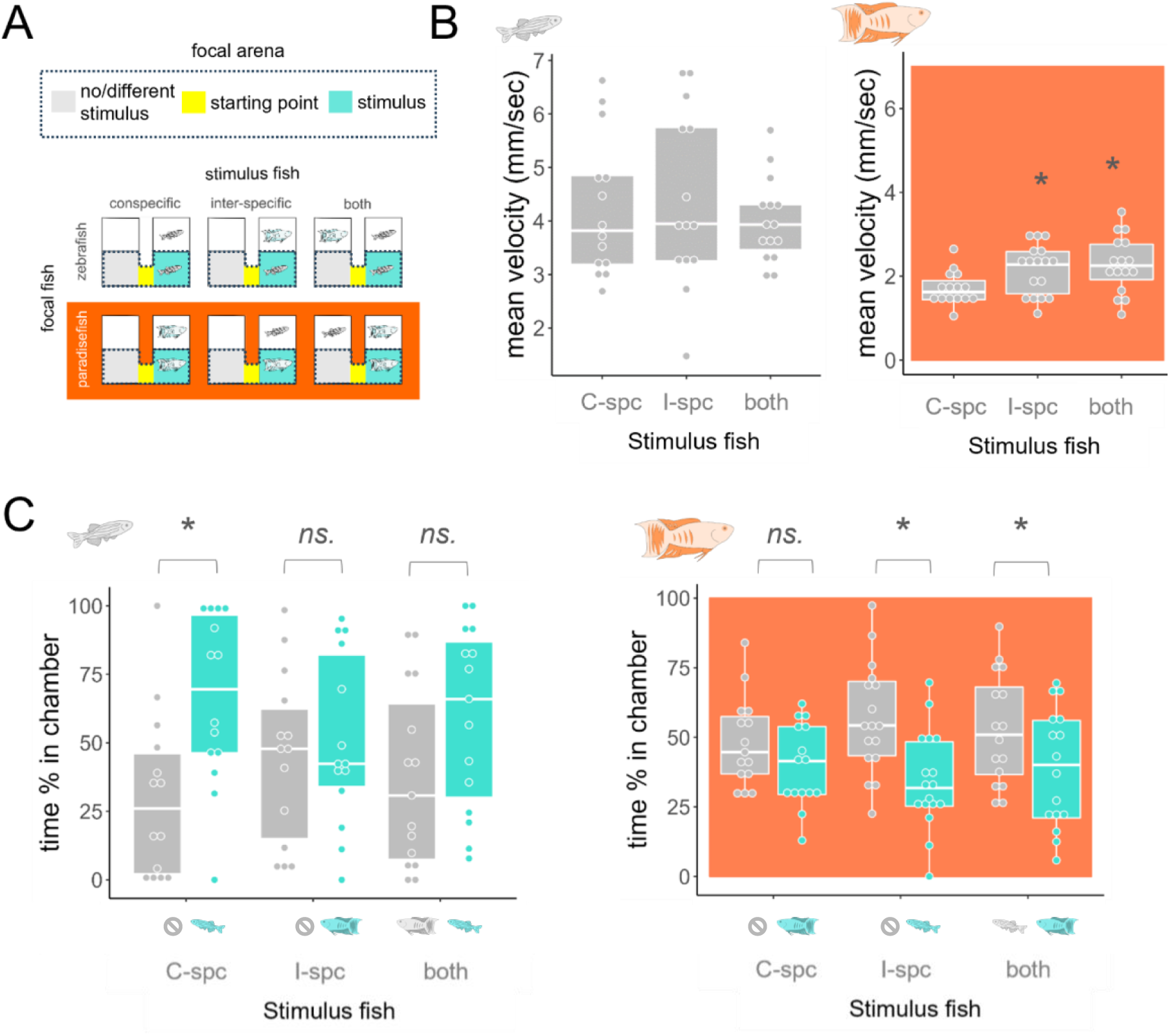
**A)** Experimental design of the intra-, inter-, and double-species challenges in which a conspecific (C-spc), an intraspecific (I-spc) or both are presented, respectively. Focal subjects were put to the starting point (yellow) of the focal arenas consisted of a no/different stimulus (grey) and a stimulus zones (turquoise). Following a fifteen minute habituation period, stimulus subjects were placed next to the stimulus zones or next to the stimulus and the no/different stimulus zones in case of the double species challenge. **B)** Mean velocity of swimming in the whole apparatus in response to each challenge presenting different stimulus fish. * represents significant difference from the swimming velocity shown in the intra-specific challenge. **C)** Percentage of time spent in the no/different stimulus (grey) or the stimulus zones (turquoise). * represents significant differences between time spent in the empty and the stimulus chambers.

While zebrafish showed similar swimming velocity in all contexts (F(2, 40)=0.369, p=0.693), paradise fish enhanced their swimming speed (F(2, 45)=4.023, p=0.0247) in inter-(estimate=0.447, SE=0.206, p=0.035) and double-species challenges (estimate=0.563, SE=0.209, p=0.009), indicating a response to the presence of the zebrafish stimulus (Figure 1B). In line with previous data, zebrafish preferred the proximity of a conspecific compared to an empty chamber (estimate=-36.17, SE=12, p=0.003) (Figure 1C left). Interestingly, the presence of a stimulus paradise fish per se did not trigger the approach of zebrafish (estimate=-6.25, SE=12, p=0.603), but was able to diminish conspecific preference in the double-species challenge (estimate=-22.31, SE=11.6, p=0.057). On the contrary, paradise fish did not snow preference towards a conspecific in neither condition (Figure 1C right): they spent approximately the same amount of time in both chambers in the intra-species challenge (estimate=8.23, SE=6.73, p=0.225), avoided zebrafish in the inter-species challenge (estimate=22.16, SE=6.32, p=0.0007), and avoided conspecifics in the double-species challenge (estimate=13.85, SE=6.52, p=0.036). In summary, in response to the presence of the other species, zebrafish have lost their original preference, while paradise fish have developed active aversion from a passive state. Interestingly, paradise fish choose the proximity of a different species over its own conspecific, but do not avoid conspecifics per se, indicating that the presence of a conspecific is not completely neutral for them but alters the response to an interspecific individual.

### Exploration of paradise fish is not biased by to the presence of a conspecific

The stress-ameliorating role of conspecifics in social species is known as social buffering, a phenomenon that has been described in a wide range of species, including zebrafish. Given the divergent responses to conspecifics in our first experiment, next, we aimed to investigate social buffering on exploration of a novel environment. One or two individuals of zebrafish or paradise fish were exposed to a 12-compartment linear maze called the slalom test (Figure 2A) and mean transition latency and overall exploration success were measured (Figure 2B and 2C). Larval zebrafish, which are known to show less discrete social activity compared to juveniles were also present as control. We found that juvenile zebrafish with a companion more actively explored the area (F(2,177)=4.709, p=0.010) than groups of larval conspecifics (estimate=0.153, SE=0.067, p=0.035) or paradise fish of the same age (estimate=0.227, SE=0.076, p=0.009), indicated by the decreased mean transition latencies (Figure 2B). Interestingly, larval and juvenile zebrafish show a bimodal exploration pattern in different conditions; alone or with a companion, respectively, which potentially indicates the presence of more than one exploration strategy in the population. Bimodality is apparent in larvae when those explore alone, while bimodality is presented in juveniles when those explore as a group of two. In contrast, paradise fish show a unimodal gaussian distribution of exploration of the novel area in every condition (Figure 2D left). All distribution types were confirmed using gaussian mixture modelling (Figure 2D right): the highest BIC values for one peak (unimodal distribution) were at assays that assessed the exploration of one juvenile zebrafish or of either one or two paradise fish. Interestingly, besides less dynamic exploration, paradise fish showed the highest success rate in exploring the whole area, irrespective of the group size (Fisher exact with simulated p-values based on 2000 replicates, p=0.0009) (Figure 2C). Based on this data the exploration of paradise fish is more effective compared to zebrafish and is not biased by the current presence of a conspecific.

**Figure 2.**
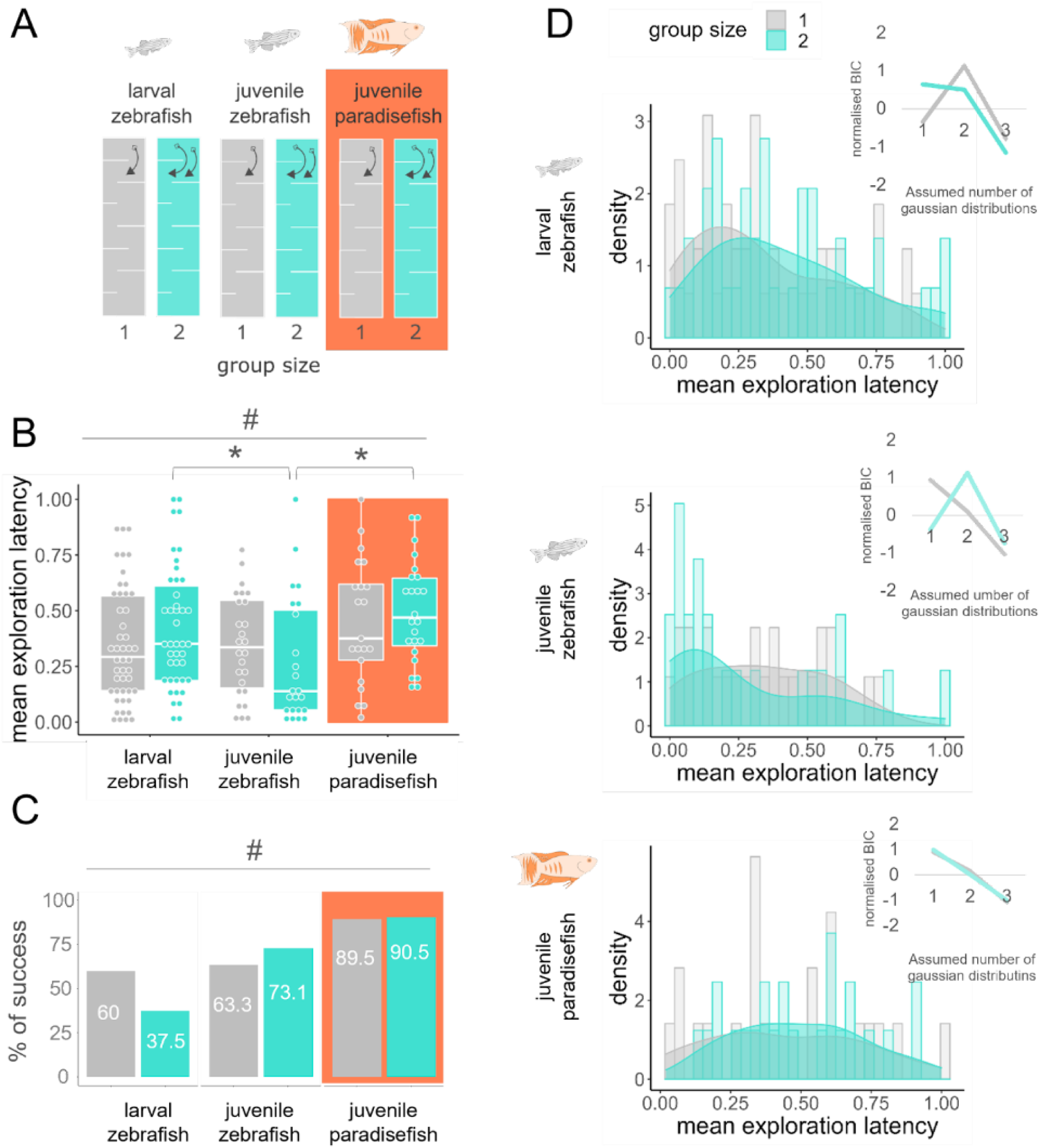
**A)** Experimental design of an exploration challenge using larval and juvenile zebrafish and juvenile paradise fish (species categories) applied alone or with a companion (group categories) to the slalom test. Colour code for the apparatuses matches the colour code of the plots. **B)** Mean transition latency (scaled and averaged entry latencies to each chamber) of subjects. # represents significant main effect of the species category, * represents significant difference between group categories. **C)** Percentage of the animals that reached the last chamber in a 10 minute long session. **D)** Distribution of exploration latencies (histograms) and the probability of their uni-, bi-, or tri-modal nature. The highest BIC values show the most likely number of independent gaussian distributions that the total distribution may consist of.

### Exploration of paradise fish is not biased by subchronic social isolation

Next, we aimed to investigate whether the subchronic (1-3 days) absence of conspecifics affects the behaviour of paradise fish. We introduced zebrafish or paradise fish to social isolation on day 0 and tested their anxiety-like responses on day 1, 2 and 3 in the swimming plus-maze (SPM) and showjump (SJ) tests (Figure 3A). Since repeated testing and the varying durations of social isolation at sampling points both bias our primary outcomes in such design, we introduced additional test-naive control groups on every sampling day. We found that socially isolated zebrafish showed increased exploration latencies (Figure 3B and 3C) (SJ: F(3,139)=4.995, p=0.003; SPM: F(3,139)=8.324, p=0.00004) and overall decreased exploration (Figure 3D and 3E) (SJ: F(3,139)=4.377, p=0.006; SPM: F(3,139)=9.096, p=0.00002) in contrast to socially housed (tested or test-naive) individuals in both tests (SJ: tested, latency: estimate=0.167, SE=0.065, p=0.017, tested, frequency: estimate=-12, SE=6.51, p=0.101, test-naive, latency: estimate=0.273, SE=0.076, p=0.001, test-naive, frequency: estimate=-24, SE=7.61, p=0.006) (SPM: tested, latency (day1/2/3): estimate=2.37/141.24/203.37, SE=52.2/52.2/52.2, p=1/0.023/0.0005, tested, frequency(day1/2/3): estimate=-0.211/-5.579/-8.947, SE=2.74/2.74/2.74, p=1/0.087/0.006, test-naive, latency(day1/2/3): estimate=-/193.92/209.59, SE=-/55.6/53, p=-/0.004/0.0005, test-naive, frequency(day1/2/3): estimate=-/-11.87/-8.737, SE=-/2.91/2.77, p=-/0.0005/0.006), an effect similar to that of chronic isolation^24^. Interestingly, despite both tests being based on the conflict between exploration of novel areas and the aversive proximity of water surface, social isolation exerted slightly different effects on their endpoints. In contrast to SJ testing, behavioural changes in the SPM test were only present from the second day for the isolated individuals (SPM group:day interactions (latency/frequency): F(2,139)=3.88/2.59, p= 0.022/0.078, see the post-hoc comparisons above), indicating that only a relatively longer period of isolation was able to modify this exploration pattern. In paradise fish, subchronic social isolation had no effect on anxiety-like behaviour in any of the tests (SJ group main effects (latency/frequency): F(3,125)=0.83/1.75, p=0.479/0.159; SPM group main effects (latency/frequency): F(3,119)= 0.49/ 3.68, p= 0.688/0.028). However, we noticed a significant decrease in exploration (frequency) in response to retesting in the species (SPM, test-naïve, frequency: estimate= -5.56,2.05,0.0231).

**Figure 3.**
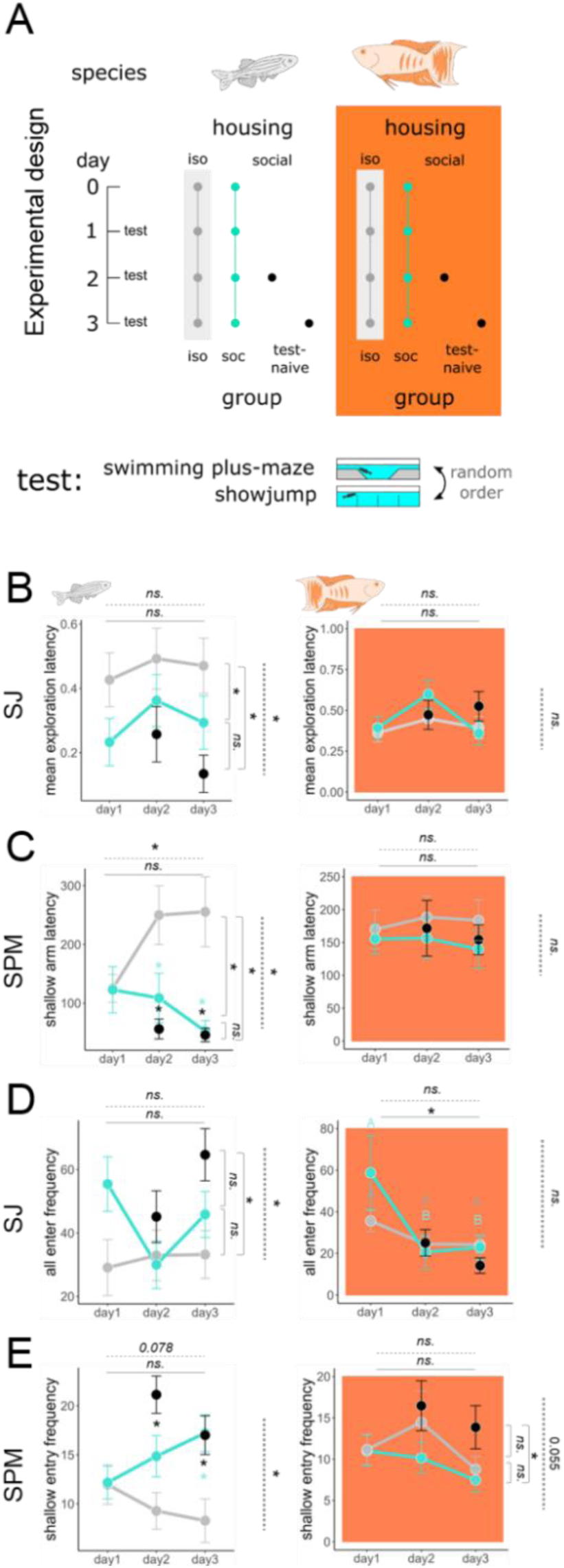
The effect of social isolation on exploration. **A) Experimental design**: Zebrafish and paradise fish were kept in isolation or in social groups for 4 consecutive days and their behavior was monitored with the SPM and SJ tests. Additional test naïve controls were tested in each testing day to control for carry-over effects of repeated testing. **B-C) exploration latencies** in the SJ (B) and the SPM (C) tests of zebrafish (left) and paradise fish (right). **D-E) exploration frequency** in the SJ (D) and the SPM (E) tests of zebrafish (left) and paradise fish (right). Horizontal solid or dashed lines with asterisk represent significant main effect of testing day or significant interaction between testing days and groups, respectively. Vertical dashed lines with asterisk represent significant main effect of the groups. Vertical braces with asterisks or different letters indicate significant post-hoc contrast between groups or testing days, respectively.

### Paradise fish and zebrafish both express anxiety-like responses

Given that a severe environmental perturbation - social isolation - did not influence the surface avoidance of paradise fish, we sought to assess whether this behaviour is associated with anxiety in this species at all. Therefore, we treated zebrafish and paradise fish with the human anxiolytic medicine buspirone and measured their avoidance in the SPM test. We also show summary measures (averages of the three sampling days) of the SPM test from our prior social isolation experiment to compare the magnitude of change in anxiety-like responses to environmental and pharmacological perturbations. We found that the same endpoint (anxiety score) that was not responsive to isolation in paradise fish (Figure 4A) (paradise fish: estimate=-0.724,SE=0.391,p=0.07), but in zebrafish (zebrafish: estimate =- 1.25, SE=0.362, p=0.001) decreased in response to buspirone in both species (paradise fish: Kruskal-wallis/posthoc Wilcoxon adjusted with FDR=p=0.009/0.014; zebrafish: Kruskal-wallis/posthoc Wilcoxon adjusted with FDR=p=0.074/0.048). Note that avoidance of zebrafish and paradise fish decreased in response to different concentrations of buspirone, 25 and 50 mg/L, respectively (Figure 4B left). Interestingly, while zebrafish enhanced their locomotion in response to the anxiolytic agent, paradise fish decreased it (Figure 4B right) (paradise fish, velocity: F(2,38)=26.884, p<0.00001; zebrafish, velocity: F(2,43)= 33.664, p= 0.006) indicating a similar dichotomy that was seen during a social challenge in our first experiment. These results indicate that surface avoidance behavior is a marker of anxiety in both species, but in contrast to zebrafish, in paradise fish it is not affected by acute social isolation.

**Figure 4.**
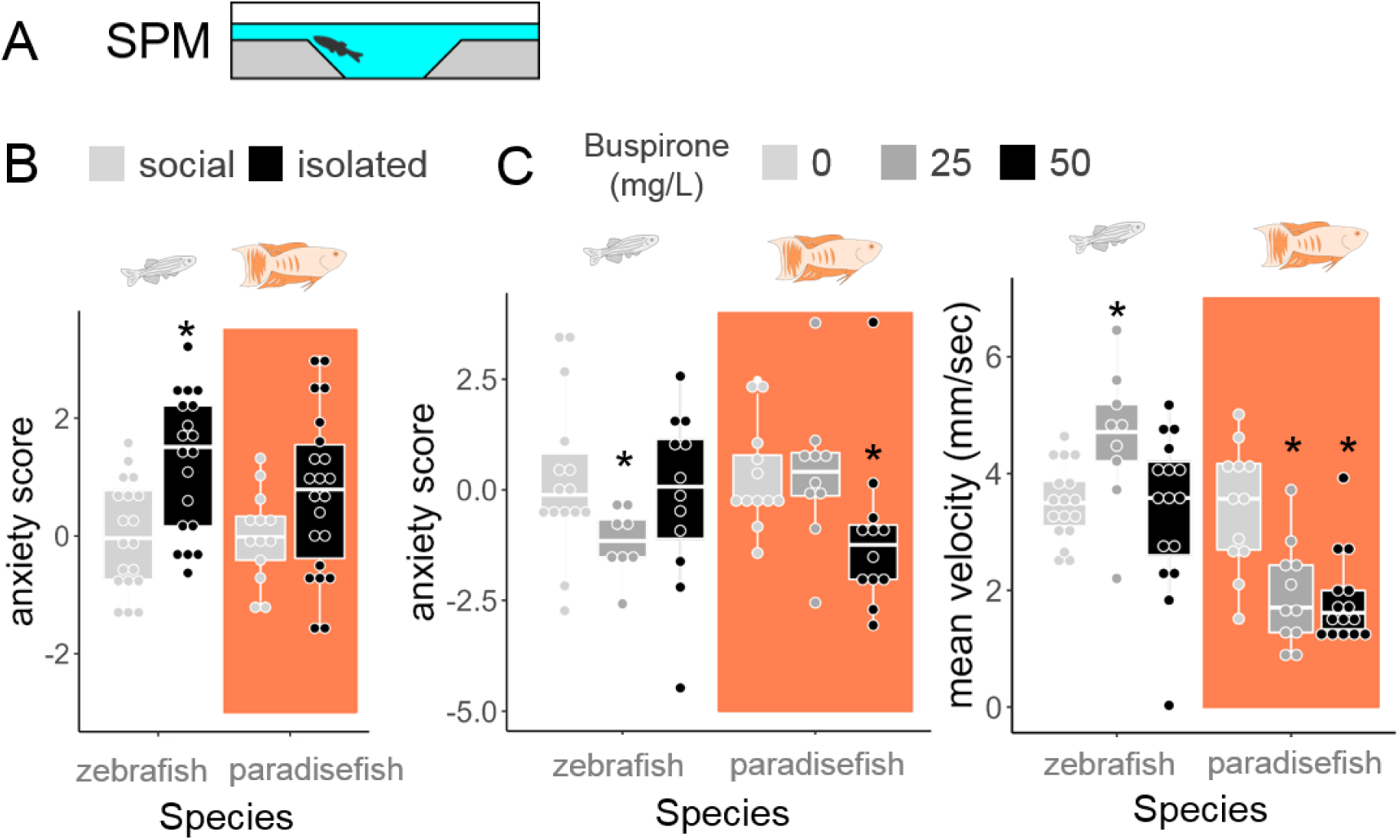
Exploration variables as markers of anxiety. A) Zebrafish and paradise fish were tested in the SPM test following social isolation stress or anxiolytic buspirone treatment. Anxiety scores were calculated from time and frequency data of shallow arm activity. B) Shallow arm activity (anxiety-score) in response to 4 days of social isolation in zebrafish (white) and paradise fish (orange). C) Shallow arm activity (anxiety-score) and mean velocity in response to buspirone in zebrafish and paradise fish. Asterisks represent significant post-hoc contrast from socially reared (social isolation) or vehicle treated (buspirone) groups following significant group main effect.

### Paradise fish show behavioural consistency through time and contexts

In our next experiment, we compared behavioural consistency in zebrafish and paradise fish. Behavioural consistency implies predictable responses through time and context which we approached with intra-test (repeatability in time) and inter-test (repeatability in contexts) correlation analysis of behavioural endpoints, respectively. In the case of inter-test correlations we used both single time point (SiM) and refined summary measurements (averages of multiple sampling events, SuM) which better cover stable, trait-like features of the individuals^38^. Animals with more consistent behaviour show stronger correlations between test-retest performances (strong repeatability) or between different test performances (strong inter-test correlations) which can be further enhanced by refined sampling (e.g. using SuMs instead of SiMs). We sampled locomotion and anxiety-like avoidance behaviour in the SPM and open tank (OT) tests in three five-minute sessions with ten-minute inter-test-intervals (ITI) (Figure 5A). We calculated between-and within-test variability (Figure 5C), bootstrap analysis-based repeatability scores (Figure 5D) and Spearman correlations (Figure 5B) of several endpoints of the two tests. Swimming velocity and the number of immobile episodes showed big within-test variability and small inter-test variability and in line with this represented the most repeatable outputs in both species. In contrast, avoidance (from the shallow arms or the center of the SPM or OT tests, respectively) showed either low within-test variability or high between-test variability, making these variables less repeatable.

**Figure 5.**
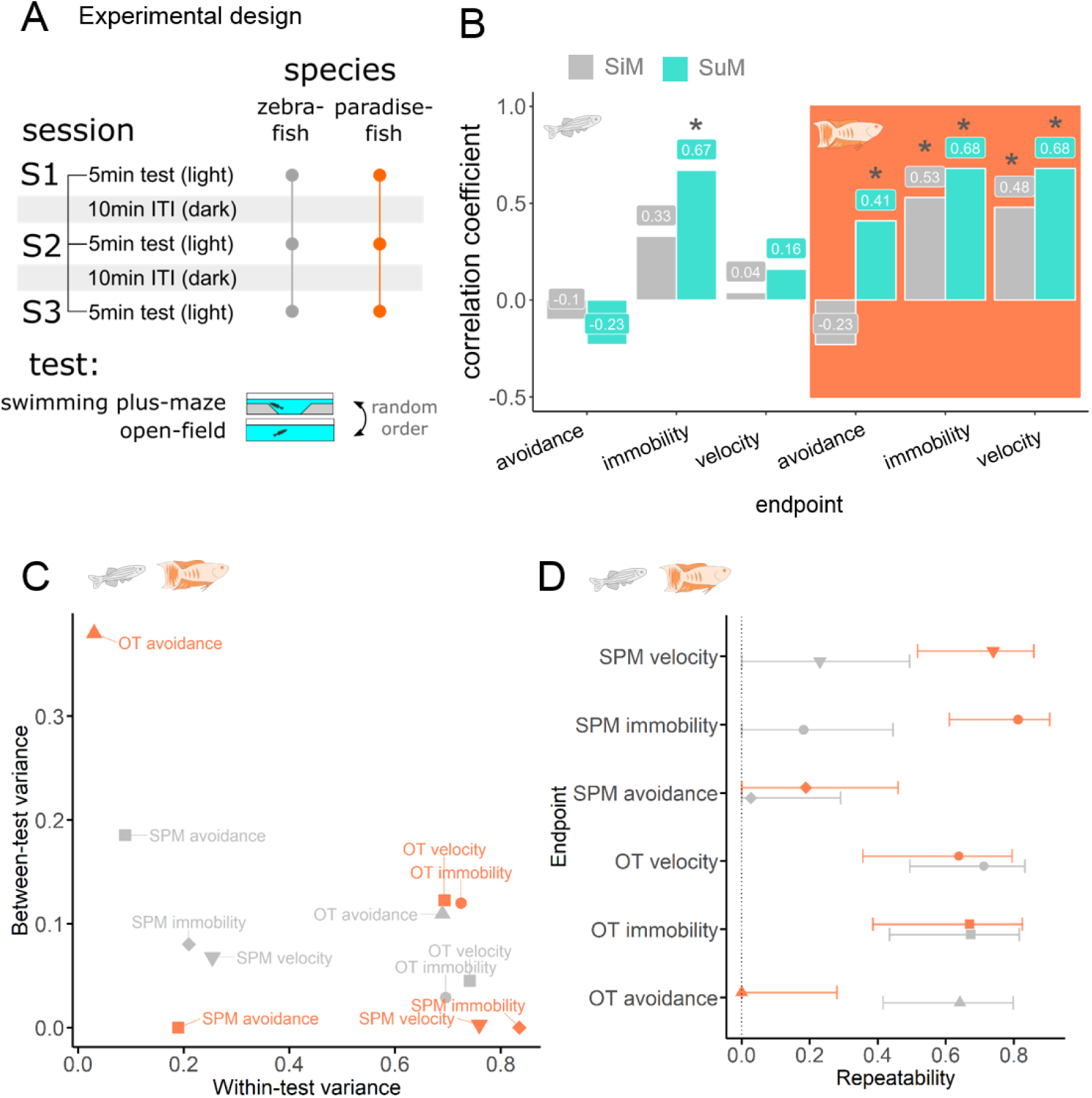
Behavioural consistency in contexts and time were investigated by conducting two type of anxiety tests (SPM and OT) in a repeated design measuring inter and intra-test variability, respectively. A) Design of the experiment. B) Inter-test correlations between similar variables measured in the SPM and OT tests. Grey bars represent correlations between single measures (one test event) and turquoise bars represent correlations between summary measures of three consecutive test events. C) Single measures of the species in the between-test-within-test variance space. D) Repeatability scores of single measures and their confidence intervals were calculated from between-and within-test variability by parametric bootstrapping. Variables with a confidence interval crossing the 0 dotted line are not considered repeatable.

In paradise fish, locomotor variables calculated either as SiMs or SuMs significantly correlated between the two tests, indicating a time-and-context-consistent locomotion (Figure 5B). In addition, using summarisation, correlations between the two tests were revealed and further enhanced using avoidance and locomotor variables, respectively. Likewise, a significant correlation between the frequency of immobile episodes in the SPM and OT emerged by the summarisation of sampling events in zebrafish. In summary, paradise fish show consistent locomotor behaviour through time and contexts and consistent avoidance behaviour through contexts. In contrast, in the case of zebrafish, behavioural consistency is limited to the expression of episodic/saccadic swimming.

### Paradise fish express different exploration strategy and level of working memory compared to zebrafish

Given these fundamental differences in exploration patterns, in our next experiment we aimed to assess by what strategy paradise fish and zebrafish explore novel areas. We used the y-maze test, an aquatic version of the rodent working memory-assessment test under the same name^*39,40*^ (Figure 6A). The idea of the rodent y-maze is that intact working memory is necessary to apply the most effective strategy in exploration, namely the coherent alternation of different arms^*39*^. Both the rate of possible actions (indirect or direct revisits, alternations) (Figure 6B) and the percent of alternations differed in the two species (Figure 6C) (action_main-effect_/species_main-effect_/action:species_interaction_: F(2,62)=302.91/15.216/82.671, p<0.001; post-hoc_(indirect revisits/direct revisits/alternations)_: estimate=11.29/0.245/-27.433, SE=2.25/2.25/2.25, p=<0.0001/0.913/<0.0001). Paradise fish show approximately 70% of alterations which exceeds the corresponding result of zebrafish and is in the range of the performance observed for mice. It is important to note that these results do not imply that zebrafish do not have working memory. Since their performance was under the random choice (50%), the results indicate that their exploration strategy is different to that of paradise fish or rodents.

**Figure 6.**
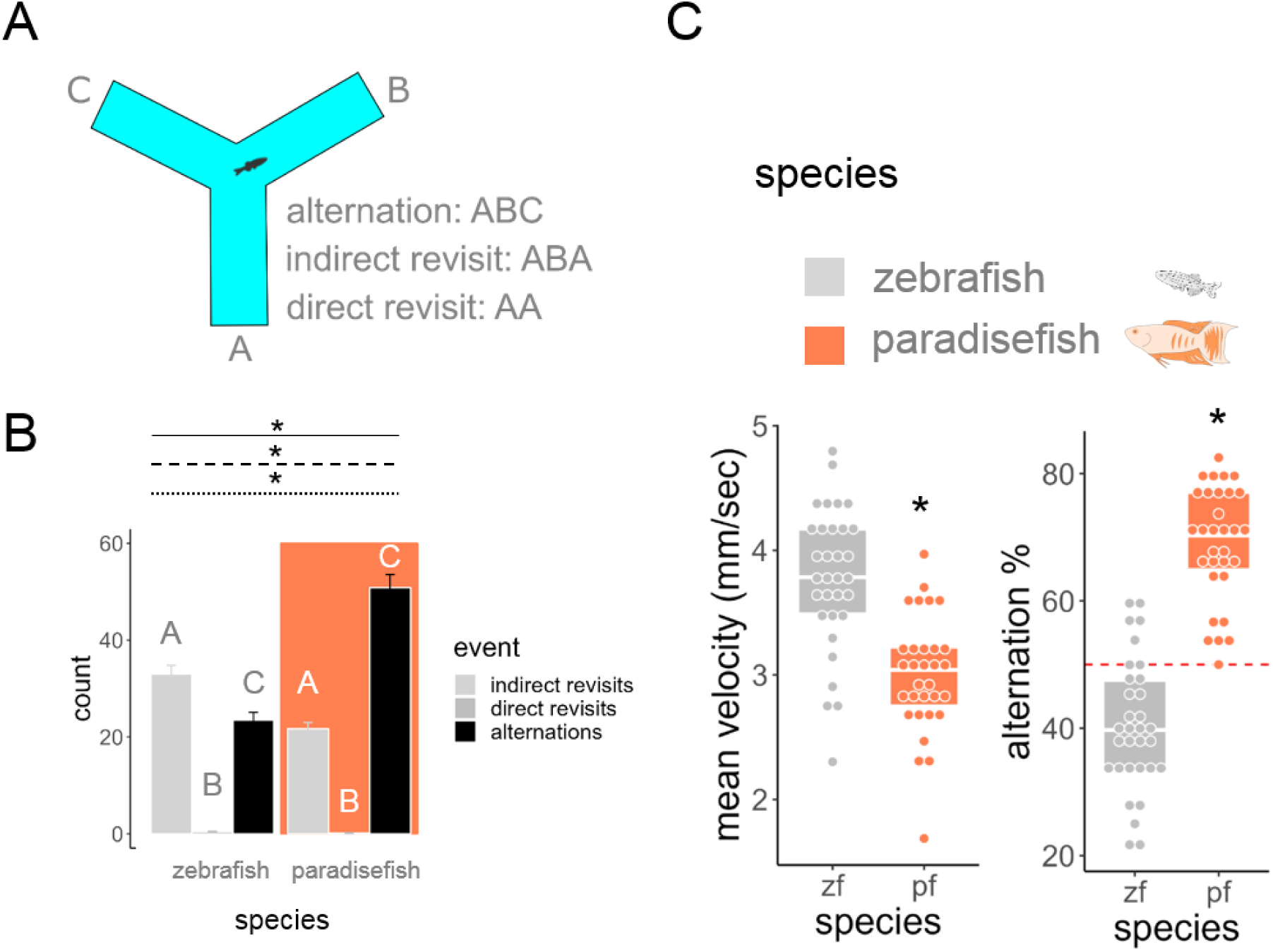
Exploration strategy of zebrafish and paradise fish. A) Schematic drawing of the y-maze platform and a list of the possible exploratory actions. B) Distribution of different exploratory actions in zebrafish (white) and paradise fish (orange). Asterisks on a solid, dashed or dotted line represent significant main effect of species, action type or significant interaction between these, respectively. Different letters represent significant post-hoc contrasts. C) mean swimming velocity and alternation % of the two species. Asterisks represent significant species differences. Red dashed line represents 50% alterations, meaning random choice in the y-maze test.

## DISCUSSION

Here we introduce the juvenile paradise fish model to improve individual-subject-based assays of exploration, anxiety and working memory. We showe that the outcome of these tests is less biased by external factors when using paradise fish compared to zebrafish. We also showe that this difference is likely due to zebrafish having a stronger social preference compared to paradise fish which can be a biasing factor either when tested alone or in groups.

Social behaviour of paradise fish was investigated before, but existing literature focuses more on territorial aggression rather than different aspects of sociability and applied adult fish instead of juveniles ^30,33,34,41^. Despite the fact that paradise fish do not show an active social approach in the U-shape sociability test or its exploratory behaviour is not affected by the absence or presence of a conspecific, some indirect effects of the social cues were apparent in our study. Juvenile paradise fish approached zebrafish individuals in the U-shape test but only in the inter-specific challenge with the concomitant presence of another paradise fish individual. These results contradict earlier data of Gerlai and Hogan who described paradise fish conspecifics as social reinforcers in an intra-specific challenge and different species as neutral in an inter-specific challenge^34^. They, however, used adult male paradise fish in their experiments and discussed the potential role of territorial aggression which is not apparent in juveniles used in the current study. Such conditional approach of a different species can be interpreted as mild “curiosity” which only worth the risk if social companions are present. Paradise fish enhanced their swimming velocity in response to the presence of a zebrafish individual while lowering it in response to the clinical anxiolytic agent buspirone. This indicates that the behaviour is anxiety-related and that paradise fish are anxious in the close proximity of a different species such as zebrafish. Approaching individuals of another species in the presence of a conspecific, however, can be considered an indication of a social buffering effect on this anxiety-like state. This is supported by the previous results of Miklósi et al who found that social buffering of the anti-predator behaviour is apparent in juvenile paradise fish^31^. The fact that subchronic social isolation does not affect the behaviour of paradise fish is also an indicator of the less pronounced social reference in this species. This also highlights the potential for using the paradise fish in experimental paradigms that involve the separation of individuals for several days. Such isolation periods present a serious confounding factor for other mainstream behavioural models such as zebrafish^24^, mice^42^ or rats^43^. In summary, while some social drive is apparent in paradise fish, the primary outcomes of the U-shape sociability test, the Slalom exploration test or the Swimming plus-maze and Showjump anxiety tests are not affected by the acute or subchronic presence or absence of a conspecifics in this species.

We also showed that paradise fish are more successful in novelty exploration using multiple paradigms. Despite the overall slower swimming velocity of this species compared to age-matched zebrafish, a significantly higher percentage of juvenile paradise fish managed to complete the slalom maze.

Paradise fish also show more consistent exploration through time and contexts, indicated by the fact that outcomes of different exploration assays are more repeatable and more correlated with each other compared to the performance of zebrafish in the same paradigms. This superior efficacy could be explained by different, though not mutually exclusive hypotheses. First, as detailed above, paradise fish are less distracted by the presence or the absence of social cues and this can be a reason for effective exploration on their own. Another explanation is that paradise fish explore novel areas using a different strategy than zebrafish. Finally paradise fish may also be less disturbed by the physical manipulation of moving them between tanks during the tests.

Using the y-maze working memory assay on both species we showed that the majority of explorative entries are indirect revisits (ABA) in the case of zebrafish and alternations (ABC) when paradise fish are tested. Using a strategy based on indirect revisits potentially indicates a more cautious home-base-like exploration, while alterations are the quickest way to explore a novel area. Previous data also suggested that paradise fish show a more efficient strategy that enables the exploration of significantly more novel compartments than simulated random walks^44^. The strategy that we described here is similar to the one shown by rodents and humans^40^. It also highlights that paradise fish are an ideal choice for reductionist high-throughput screening that is more translatable to mammalian species and ultimately to humans.

The y-maze can also be used to measure the working memory of zebrafish, albeit in a more indirect way^45^. Speculating about the effectiveness of these alternative strategies on exploration, one could argue that zebrafish can also be successful in this paradigm by compensating for its slower exploration strategy with higher swimming speed. It needs to be highlighted, however, that efforts to understand the explorative strategy of an R-strategist^46^ shoal fish evaluated alone are deemed to be highly speculative. It is likely that the most successful strategy for larval zebrafish that find themselves alone is to return to their shoal. Based on our results, therefore, we suggest that paradise fish are potentially more useful for translational research than zebrafish, as their exploratory behavior, tested in individual settings is more straightforward to interpret.

The use of larval and juvenile zebrafish in assays presenting either social or non-social novelty challenges are dominating the field of reductionist neuroscience. The strong, species-specific social drive, however, makes it hard to interpret their behaviour in non-social contexts. Paradise fish, on the other hand, possess all the features that made zebrafish a predominant model for neuroscience and also show unbiased challenge-coping behaviour. In addition, important pre-requirements of a useful model were fulfilled by the current description of the paradise fish reference genome^28^ and efficient husbandry^29^. Figure 7 summarises the outcomes of the different assays, categorising them as social and non-social novelty challenges in juvenile zebrafish and paradise fish. Our results show that zebrafish is a better choice for social assays, whereas paradise fish are superior models for the non-social novelty tests. We believe that our results provide compelling evidence for the use of the paradise fish as a complementary model in biomedical sciences, that could offer further and potentially more translatable insights to the background mechanisms of affective and cognitive processes.

**Figure 7.**
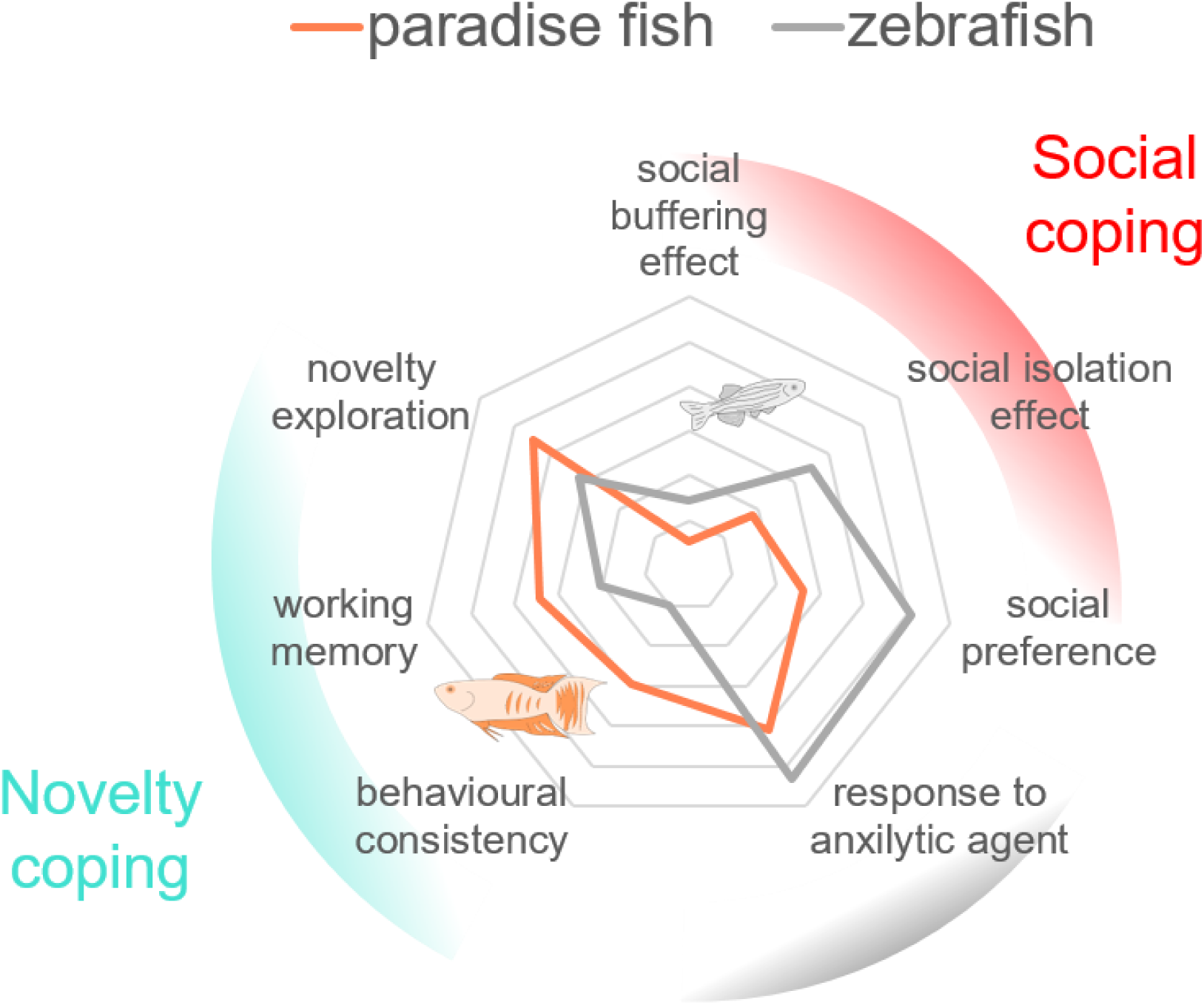
Effect sizes of comparison between zebrafish and paradise fish in several assays tracing out the possible fields of use of the two species.

## MATERIALS AND METHODS

### Animals

Wild-type zebrafish (AB) and paradise fish lines were maintained in the animal facility of ELTE Eötvös Loránd University according to standard protocols^29,47^. Experimental subjects were male and female animals aged between 10 and 30 days post fertilization (dpf). Fish were maintained in a standard 14 h/10 h light/dark cycle. Feeding of zebrafish and paradise fish larvae started at 5 dpf with commercially available dry food (a 1:1 combination of <100 µm and 100-200µm Zebrafeed, Sparos). After 15 dpf, juvenile fish were fed using dry food with gradually increasing particle size (200-400µm Zebrafeed, Sparos) combined with fresh brine shrimp hatched in the facility. Animals were anaesthetised with lidocaine and terminated with tricaine immediately after each experiment. All protocols used in our study were approved by the Hungarian National Food Chain Safety Office (Permits #PEI/001/1458-10/2015, #PE/EA/2483-6/2016 and #PE/EA/406-7/2020).

### Environmental modifications

Sub-chronic social isolation of subjects was conducted between 27 and 30 dpf. Animals originating from the same spawning were randomly allocated to social rearing or social isolation for both species. Isolated animals were kept in white opaque plastic tanks (52×35×46 mm, depth x width x length) depriving the individuals of sensory cues from conspecifics^24^. Tanks were filled with fish system water, half of the volume of which was replaced on a daily basis. Socially reared control animals were subjected to similar conditions, with the exception of the size of their aquarium, which was matched to the greater number of larvae, providing ∼50 ml volume for each individual.

### Drug treatments

Buspirone (Sigma Aldrich, Cat. no. 33386-08-2) was dissolved in system water and administered as a water bath for 10 min followed by a 5 min washout in compartments of a 12-well plate followed by a behavioural test immediately. Each compartment contained 1.5 ml of treatment solution. We applied 0 (vehicle), 25 and 50 mg/L concentrations, which were selected based on our previous studies and pilot experiments^24,48^.

### Behavioural procedures

#### Sociability test

To measure exploration in social context the sociability test of the Dreosti laboratory^19^ and the slalom test developed by our group (see below) were used. The social preference test apparatus is a U-shaped platform (40x32 mm) consisting of two identical arms, with glass window partitions enabling only visual communication between the fish (Figure 1A). The test is based on the visually-guided preference of zebrafish toward conspecifics. A fifteen minute habituation period followed by a fifteen minute challenge period, without or with the presence of a stimulus animal, respectively, according to the protocol from the original paper and confirmed by our previous studies^24^. Time spent in each zone was measured.

#### Slalom test

The slalom test was recently developed by our laboratory to measure the the exploratory drive of larval or juvenile fish, particularly in a social context. The relatively long test apparatus (15×20×85 mm, depth x width x length) consists of 12 equal-sized chambers, each visible only from the adjacent ones (Figure 2A). This way the test offers enough novelty for the fish and separates the population into individuals who did and individuals who did not succeed to reach the last chamber. One or two individuals per maze were placed in the first chamber then their behaviour was recorded for ten minutes. Enter latencies (sec) to each chamber (2^nd^, 3^rd^, …12^th^) were scaled for the investigated population from 0 to 1 then these were averaged between chambers resulting in the variable “mean transition latency” (0-1). Percent of success means the proportion of the group that successfully reached the 12^th^ chamber. Sessions were analysed up to 10 minutes, or until the (last) individual reached the last chamber.

#### Swimming plus-maze test

To measure avoidance responses to novelty the swimming plus-maze (SPM), the showjump (SJ) and the open tank (OT) tests were used. The SPM test from our laboratory (Varga et al., 2018) is a +-shaped platform consisting of two plus two opposite arms, different in depth, connected by a center zone (Figure 3A). Larval and juvenile zebrafish prefer the deep over the shallow arms and the center zone, a preference linked to anxiety-like motivational states. Ten-minute-long (experiment 3 and 4) and three five-minute-long (experiment 5) tests were conducted. Time spent in every zone, the latency to enter each zone and the mean of overall velocity were measured. Standard anxiety scores were calculated as scaled time minus scaled latency multiplied with -1, representing a positively correlated measure of anxiety-like behaviour in these tests. For the validation procedure and detailed specification of the test, see Varga et al^48^.

#### Show jump test

The SJ test (15×20×42 mm, depth x width x length) consists of 4 equal-sized chambers separated by 10 mm high walls (Figure 3A). Subjects need to swim above the wall, close to the water surface to enter a novel chamber. The behaviour in the test, similarly to the behaviour in the SPM, is based on the conflict between explorative motivation and novelty-induced anxiety-like state. Ten-minute-long tests were conducted. The number of all chamber entries and mean trasition latency were measured similarly as in the case of the slalom test.

#### Open tank test

OT tests were conducted in 6-well-plates. The behaviour in the test is based on the natural aversion of highly exposed, open areas. An inner circle that covers eighty percent of the total volume was traced out representing the aversive central area. Time spent in the periphery, the number of immobile episodes, and overall mean velocity were measured.

#### Y-maze test

To characterise exploration strategies and measure working memory the y-maze test was used. The aquatic y-maze consists of three equal-sized arms (each 5×7.5×30 mm, depth x width x length) (Figure 6A). Ten-minute-long tests were conducted. We recorded alternations (three consecutive visits are done to different chambers), direct revisits (two consecutive visits are done to the same chamber) and indirect revisits (following two consecutive visits to different chambers the third is done to the first chamber). The sequences of these events were calculated in a sliding window, meaning that following the third visit every subsequent visit defined a new sequence.

### Experimental design and analysis

In **Experiment 1** we aimed to compare the sociability of the paradise fish and zebrafish in different conditions. We observed either 30 dpf paradise fish (n=15, 17, 16) or zebrafish (n=14, 14, 15) as they faced a conspecific (intraspecies challenge), an individual of the other species (interspecific challenge) or individuals of both species (double challenge) in the U-shape sociability test (Figure 1A). The intraspecific challenge is an indicator of the social activity of each species in baseline conditions while the double challenge measures the same during a mild perturbance, i.e., the presence of the other (non-predatory) species. Intraspecific challenges are controls to determine whether changes in the other conditions are species-specific e.g., triggered by a conspecific. Stimulus animals were naïve to the test and were previously housed in different tanks. The behaviour of the subjects was video recorded.

In **Experiment 2** we aimed to compare the exploratory drive in social and non-social contexts in paradise fish and zebrafish. We applied either 30 dpf paradise fish (n=21, 19) or 8 (n=24, 30) or 30 dpf zebrafish (n=26, 30) to the slalom test individually or accompanied by a conspecific (Figure 2A). The behavior was recorded for 10 minutes.

In **Experiment 3** we aimed to compare the effect of sub-chronic social isolation on anxiety-like states of paradise fish and zebrafish. We allocated paradise fish (n=13, 23) or zebrafish (n=19, 19) to social or isolated housing at day 0 (27 dpf), then assessed their behaviour from day 1 to 3 in the SPM and SJ tests for 10 minutes each day. To control for a carry-over effect of repeated testing we applied additional test naive paradise fish (n=13, 14) and zebrafish (n=15, 18) individuals from day 2 to 3 (Figure 3A). The behaviour was video recorded.

In **Experiment 4** we aimed to validate whether surface avoidance behaviour can be considered as a marker of anxiety-like states in paradise fish. We treated either 30 dpf paradise fish (n=12, 12, 15) or zebrafish (n=17, 9, 16) with different concentrations of the clinical anxiolytic buspirone and measured their surface avoidance behaviour in the SPM.

In **Experiment 5** we aimed to compare behavioural consistency through time and contexts of paradise fish and zebrafish. We applied either paradise fish (n=24) or zebrafish (n=24) to three sessions of OT then three sessions of SPM for five minutes each, interrupted by 10 minutes inter-session-intervals (Figure 5A).

In **Experiment 6** we aimed to assess the exploration strategies and compare the working memory of paradise fish and zebrafish. We applied either 30 dpf paradise fish (n=31) or zebrafish (n=32) to the aquatic version of a y-maze test for 10 minutes.

The behaviour of the subjects were recorded in a Zantiks MWP unit (Zantiks Ltd., Cambridge, UK) and analysed with Noldus Ethovision XT^49^ in each experiment.

### Statistical analysis

Statistical analysis was done in the R statistical environment^50^. Hypothesis testing on multiple group design experiments was done using linear models (ANOVA or Kruskal-Wallis rank sum test) followed by post-hoc contrasts (t-test or Wilcoxon test) depending on the distribution of data, adjusted with false discovery rate approach^51^. Hypothesis testing on single treatment group design experiments was done using Student’s *t*-tests.

Correlations were calculated using the Pearson method with the *rcorr* package. Repeatability analysis was done according to the analysis pipeline of Answer et al.^52^ for each test-type by calculating the proportion of within-test variance out of total variance, as shown below: within test variance/(within test + between test variance), where within-test (or between-individual) variance means the variation of the group’s behaviour within a given test, and between-test (or within-individual) variance means variation of individuals’ behaviour across weeks (test repetitions) for a given test-type. The R package *rptR* (version 0.9.22) was used to calculate variances and repeatability estimates similarly to previous investigations. The function calculates 95% confidence intervals of estimates by parametric bootstrapping, with the number of parametric bootstraps for interval estimation set to 1000. Estimates with confidence intervals that did not include 0 were considered statistically significant. After examining normality of variables, the variances, adjusted repeatability estimates and their uncertainty were calculated for all behavioural variables (avoidance, immobility, velocity). Gaussian mixture modeling was done using the R package *mclust* on the mean transtition latency variables. The modality of distribution was decided based on the highest bayesian information criterion (BIC) value. BIC values were scaled to 0-1 in each group and ploted together to show the differences in distribution types between species of particular age groups.

## CONFLICT OF INTEREST

The authors declare no conflict of interest.

## AUTHOR CONTRIBUTION

Conceptualization: ZKV, DP, EM, AM, MV.

Funding acquisition: MV, AM.

Investigation: ZKV, DP, TC.

Methodology: ZKV.

Writing – original and revised text: ZKV, EM, AM, MV.

## ACKNOWLEDGEMENT

We would like to thank Anita Rácz for fish care. This work was supported by the ELTE Eötvös Loránd University Institutional Excellence Program Grant 1783-3/2018/FEKUTSRAT. MV is a János Bolyai fellow of the Hungarian Academy of Sciences.

## Notes

### Competing Interest Statement

The authors have declared no competing interest.

### Summary of Updates

We have added the Conflict of Interest, Author Contribution and Acknowledgement sections that were missing from the original manuscript.

## REFERENCES

1. Chen, Z.-Y. & Zhang, Y. Animal models of Alzheimer’s disease: Applications, evaluation, and perspectives. Zool. Res. 43, 1026–1040 (2022).

2. Negi, S., Kumar, S. & Singh, A. Preclinical In Vivo Drug Development Studies: Limitations, Model Organisms, and Techniques. in Drugs and a Methodological Compendium : From bench to bedside (eds. Rajput, V. S. & Runthala, A.) 149–171 (Springer Nature, Singapore, 2023). doi:10.1007/978-981-19-7952-1_6.

3. Johan Arief, M. F., Choo, B. K. M., Yap, J. L., Kumari, Y. & Shaikh, M. F. A Systematic Review on Non-mammalian Models in Epilepsy Research. Front. Pharmacol. 9, 387505 (2018).

4. Braems, E., Tziortzouda, P. & Van Den Bosch, L. Exploring the alternative: Fish, flies and worms as preclinical models for ALS. Neurosci. Lett. 759, 136041 (2021).

5. Anderson, D. J. & Adolphs, R. A Framework for Studying Emotions Across Phylogeny. Cell 157, 187–200 (2014).

6. Harris, M. P., Henke, K., Hawkins, M. B. & Witten, P. E. Fish is Fish: the use of experimental model species to reveal causes of skeletal diversity in evolution and disease. J. Appl. Ichthyol. 30, 616–629 (2014).

7. Gerlai, R. Fish in behavior research: Unique tools with a great promise! J. Neurosci. Methods 234, 54–58 (2014).

8. Schartl, M. Beyond the zebrafish: diverse fish species for modeling human disease. Dis. Model. Mech. 7, 181–192 (2014).

9. Szabó, N. et al. The paradise fish, an advanced animal model for behavioral genetics and evolutionary developmental biology. J. Exp. Zoolog. B Mol. Dev. Evol. (2023) doi:10.1002/jez.b.23223.

10. Choi, T.-Y., Choi, T.-I., Lee, Y.-R., Choe, S.-K. & Kim, C.-H. Zebrafish as an animal model for biomedical research. Exp. Mol. Med. 53, 310–317 (2021).

11. Kalueff, A. V., Stewart, A. M. & Gerlai, R. Zebrafish as an emerging model for studying complex brain disorders. Trends Pharmacol. Sci. 35, 63–75 (2014).

12. Lovett-Barron, M. et al. Ancestral Circuits for the Coordinated Modulation of Brain State. Cell 171, 1411-1423.e17 (2017).

13. Wolman, M. A., Jain, R. A., Liss, L. & Granato, M. Chemical modulation of memory formation in larval zebrafish. Proc. Natl. Acad. Sci. 108, 15468–15473 (2011).

14. Leung, L. C., Wang, G. X. & Mourrain, P. Imaging zebrafish neural circuitry from whole brain to synapse. Front. Neural Circuits 7, (2013).

15. Leung, L. C., Wang, G. X. & Mourrain, P. Imaging zebrafish neural circuitry from whole brain to synapse. Front. Neural Circuits 7, (2013).

16. Arunachalam, M., Raja, M., Vijayakumar, C., Malaiammal, P. & Mayden, R. L. Natural History of Zebrafish (Danio rerio) in India. Zebrafish 10, 1–14 (2013).

17. Lee, C. J., Paull, G. C. & Tyler, C. R. Improving zebrafish laboratory welfare and scientific research through understanding their natural history. Biol. Rev. 97, 1038–1056 (2022).

18. White, L. J., Thomson, J. S., Pounder, K. C., Coleman, R. C. & Sneddon, L. U. The impact of social context on behaviour and the recovery from welfare challenges in zebrafish, Danio rerio. Anim. Behav. 132, 189–199 (2017).

19. Dreosti, E., Lopes, G., Kampff, A. & Wilson, S. Development of social behavior in young zebrafish. Front. Neural Circuits 9, (2015).

20. Tunbak, H., Vazquez-Prada, M., Ryan, T. M., Kampff, A. R. & Dreosti, E. Whole-brain mapping of socially isolated zebrafish reveals that lonely fish are not loners. eLife 9, e55863 (2020).

21. Fontana, B. D. et al. Using zebrafish (Danio rerio) models to understand the critical role of social interactions in mental health and wellbeing. Prog. Neurobiol. 208, 101993 (2022).

22. Oliveira, R. F. & Faustino, A. I. Social information use in threat perception: Social buffering, contagion and facilitation of alarm responses. Commun. Integr. Biol. 10, e1325049 (2017).

23. Faustino, A. I., Tacão-Monteiro, A. & Oliveira, R. F. Mechanisms of social buffering of fear in zebrafish. Sci. Rep. 7, 44329 (2017).

24. Varga, Z. K. et al. Conserved Serotonergic Background of Experience-Dependent Behavioral Responsiveness in Zebrafish (Danio rerio). J. Neurosci. 40, 4551–4564 (2020).

25. Forsatkar, M. N., Safari, O. & Boiti, C. Effects of social isolation on growth, stress response, and immunity of zebrafish. Acta Ethologica 20, 255–261 (2017).

26. Shams, S., Chatterjee, D. & Gerlai, R. Chronic social isolation affects thigmotaxis and whole-brain serotonin levels in adult zebrafish. Behav. Brain Res. 292, 283–287 (2015).

27. Shams, S., Amlani, S., Buske, C., Chatterjee, D. & Gerlai, R. Developmental social isolation affects adult behavior, social interaction, and dopamine metabolite levels in zebrafish. Dev. Psychobiol. 60, 43–56 (2018).

28. Fodor, E. et al. The reference genome of the paradise fish (Macropodus opercularis). BioRxiv Prepr. Serv. Biol. 2023.08.10.552018 (2023) doi:10.1101/2023.08.10.552018.

29. A, R. et al. Housing, Husbandry and Welfare of a ‘Classic’ Fish Model, the Paradise Fish (Macropodus opercularis). Anim. Open Access J. MDPI 11, (2021).

30. Csányi, V., Tóth, P., Altbäcker, V., Dóka, A. & Gervai, J. Behavioural elements of the paradise fish (Macropodus opercularis). I. Regularities of defensive behaviour. Acta Biol. Hung. 36, 93–114 (1985).

31. Miklósi, A., Pongrácz, P. & Csányi, V. The ontogeny of antipredator behavior in paradise fish larvae (Macropodus opercularis) IV. The effect of exposure to siblings. Dev. Psychobiol. 30, 283–291 (1997).

32. A, M., V, C. & R, G. Antipredator behavior in paradise fish (Macropodus opercularis) larvae: the role of genetic factors and paternal influence. Behav. Genet. 27, (1997).

33. Miklósi, A., Haller, J. & Csányi, V. The Influence of Opponent-Related and Outcome-Related Memory on Repeated Aggressive Encounters in the Paradise Fish (Macropodus opercularis). Biol. Bull. 188, 83–88 (1995).

34. R, G. & Ja, H. Learning to find the opponent: an ethological analysis of the behavior of paradise fish (Macropodus opercularis) in intra- and interspecific encounters. J. Comp. Psychol. Wash. DC 1983 106, (1992).

35. Csányi, V., Csizmadia, G. & Miklosi, A. Long-term memory and recognition of another species in the paradise fish. Anim. Behav. 37, 908–911 (1989).

36. Warren, J. M. Reversal learning by paradise fish (Macropodus opercularis). J. Comp. Physiol. Psychol. 53, 376–378 (1960).

37. Dreosti, E., Lopes, G., Kampff, A. R. & Wilson, S. W. Development of social behavior in young zebrafish. Front. Neural Circuits 9, (2015).

38. Varga, Z. K. et al. Improving anxiety research: novel approach to reveal trait anxiety through summary measures of multiple states. 2023.06.01.543235 Preprint at 10.1101/2023.06.01.543235 (2023).

39. Kraeuter, A.-K., Guest, P. C. & Sarnyai, Z. The Y-Maze for Assessment of Spatial Working and Reference Memory in Mice. in Pre-Clinical Models 105–111 (Humana Press, New York, NY, 2019). doi:10.1007/978-1-4939-8994-2_10.

40. Cleal, M. et al. The Free-movement pattern Y-maze: A cross-species measure of working memory and executive function. Behav. Res. Methods 53, 536–557 (2021).

41. Csányi, V., Tóth, P., Altbäcker, V., Dóka, A. & Gervai, J. Behavioural elements of the paradise fish (Macropodus opercularis). II. A functional analysis. Acta Biol. Hung. 36, 115–130 (1985).

42. Biro, L. et al. Post-weaning social isolation in male mice leads to abnormal aggression and disrupted network organization in the prefrontal cortex: Contribution of parvalbumin interneurons with or without perineuronal nets. Neurobiol. Stress 25, 100546 (2023).

43. Toth, M., Mikics, E., Tulogdi, A., Aliczki, M. & Haller, J. Post-weaning social isolation induces abnormal forms of aggression in conjunction with increased glucocorticoid and autonomic stress responses. Horm. Behav. 60, 28–36 (2011).

44. Székely, S., Havas, I. & Csányi, V. How paradise fish (Macropodus opercularis, L.) explores a chessboard. Acta Biol. Acad. Sci. Hung. 29, 401–406 (1978).

45. Cleal, M. & Parker, M. O. Moderate developmental alcohol exposure reduces repetitive alternation in a zebrafish model of fetal alcohol spectrum disorders. Neurotoxicol. Teratol. 70, 1–9 (2018).

46. MacArthur, R. & Wilson, E. The Theory of Island Biogeography. (Princeton university press, 2001).

47. Westerfield, M. The Zebrafish Book: A Guide for the Laboratory Use of Zebrafish. (University of Oregon Press, 2000).

48. Varga, Z. K. et al. The swimming plus-maze test: a novel high-throughput model for assessment of anxiety-related behaviour in larval and juvenile zebrafish (Danio rerio). Sci. Rep. 8, 16590 (2018).

49. Noldus, L. P. J. J., Spink, A. J. & Tegelenbosch, R. A. J. EthoVision: A versatile video tracking system for automation of behavioral experiments. Behav. Res. Methods Instrum. Comput. 33, 398–414 (2001).

50. R Core Team. R: A language and environment for statistical computing. R Foundation for Statistical Computing, Vienna, Austria. (2019).

51. Benjamini, Y. & Hochberg, Y. Controlling the False Discovery Rate: A Practical and Powerful Approach to Multiple Testing. J. R. Stat. Soc. Ser. B Methodol. 57, 289–300 (1995).

52. Anwer, H. et al. An efficient new assay for measuring zebrafish anxiety: Tall tanks that better characterize between-individual differences. J. Neurosci. Methods 356, 109138 (2021).

